# Neonatal morphine and HIV synergy induce persistent neuroimmune and anxiety-related transcriptional states

**DOI:** 10.64898/2025.12.01.691653

**Authors:** Junyi Tao, Danielle Antoine, Sabita Roy

## Abstract

**Background:** Early-life opioid exposure can disrupt neurodevelopment and heighten vulnerability to anxiety and affective disorders, particularly in individuals with HIV.

**Methods:** Using single-cell RNA sequencing (scRNA-seq), we profiled adolescent brains from wild-type and HIV-1 transgenic (Tg26) mice exposed to morphine during postnatal days 2–7.

**Results:** Morphine exposure in Tg26 mice resulted in a highly dysregulated microglial phenotype, characterized by the ectopic upregulation of genes encoding neuronpeptides (*Avp, Hcrt*, and *Pmch*), while simultaneously showing a reduction in both inflammatory and homeostatic markers (*Map3k6, Lgals3, Ccl3*). Microglia also showed enhanced expression of dynorphin (*Pdyn*) and κ-opioid receptor (*Oprk1*) signaling modules implicated in dysphoria and stress-induced negative effects. Furthermore, transcriptomic mapping revealed cell-type-specific neuronal adaptations: cholinergic neurons upregulated genes linked to anxiety and arousal (*Avp, Oxt*), GABAergic neurons upregulated genes linked to condition and aversive behavior, whereas glutamatergic neurons enriched for transcripts associated with thigmotaxis and fear behaviors.

**Conclusions:** Together, these findings demonstrate that brief neonatal morphine exposure in an HIV-inflamed milieu induces persistent, cell-type-specific neuroimmune and neurotransmitter reprogramming that engages the dynorphin–KOR pathway and predisposes to anxiety- and aversion-related behaviors.

## Introduction

Opioid exposure during early development has been increasingly recognized as a critical risk factor for long-term neuropsychiatric disorders, including anxiety, stress hypersensitivity, and substance-use vulnerability (1–4). In people with HIV (PWH), opioid use disorder (OUD) further exacerbates neuroinflammation, microglial activation, and synaptic dysregulation pathways already compromised by chronic HIV infection (5–8). Although repeated or chronic opioid exposure has been shown to alter emotion-related circuitry, the impact of brief neonatal morphine exposure (NME) in the context of HIV-associated neuroinflammation on adolescent brain development remains poorly understood.

The HIV-1 transgenic (Tg26) mouse provides a tractable model of chronic viral protein mediated immune activation (9,10). We hypothesized that early morphine exposure during the postnatal period (P2–P7), a window corresponding to the late third-trimester human brain, would produce persistent, cell-specific transcriptional reprogramming of glial and neuronal populations that predisposes to anxiety-like behavior. To test this, we performed single-cell RNA sequencing of adolescent brains from wild-type (WT) and Tg26 mice exposed to morphine neonatally. By resolving cell-type-specific trajectories, we aimed to identify how HIV-related neuroinflammation amplifies morphine-induced transcriptional changes in microglia and discrete neuronal subtypes implicated in anxiety and aversion.

## Materials and methods

### Neonatal Morphine exposure (NME) paradigm

HIV-1 transgenic (Tg26) and C57BL/6J (wild-type, WT) female mice aged 10–12 weeks were paired with healthy males of the same age and genetic background. Females were monitored daily for pregnancy and were individually housed upon confirmation of gestation. After delivery, pups from both Tg26 and WT litters were exposed to morphine beginning on postnatal days 6–7 and continuing for a total of five consecutive days. Morphine was administered subcutaneously at 5 mg/kg/day, a dose previously validated in our NME model to produce reliable analgesic effects while minimizing excessive opioid exposure. Following the final morphine injection, pups remained with their dams until weaning at postnatal day 21. After weaning, mice were sexed, group-housed according to institutional guidelines, and maintained until midbrain tissue collection during adolescence (4–5 weeks of age). All procedures were conducted with prior approval from the University of Miami Institutional Animal Care and Use Committee (IACUC) and adhered to the National Institutes of Health Guide for the Care and Use of Laboratory Animals.

### 10× genomics single-cell RNA sequencing

10x Genomics Chromium Fixed RNA Profiling was performed by the Hussman Institute for Human Genomics (HIHG) at the University of Miami. Cell suspensions were assessed using the Nexcelom Cellometer K2. The suspensions were prepped following the Chromium Fixed RNA Profiling Reagent Kits for Multiplexed Samples User Guide. Gene expression libraries were sequenced by Illumina NovaSeq X Plus.

### scRNA-seq analysis

Raw fastq files were initially processed with Cell Ranger multi pipeline (v.8.0.1, 10X Genomics) with the Chromium Mouse Transcriptome Probe Set v1.0.1 and default parameters. Processed count matrix were further processed with the Seurat package (v.5.0)(11). Cell type were annotated with sc-type package(12). Differentially expressed genes (DEGs) between groups for all cell types were identified using the FindMarkers function in Seurat, employing a built-in Wilcoxon test with Benjamini-Hochberg correction of p-values. Genes with an absolute Log2 fold change (|Log2FC|) > 0.4 and an adjusted p-value < 0.01 were considered statistically significant. QIAGEN Ingenuity Pathway Analysis (IPA) were used for canonical pathway analysis and downstream biological and disease function analysis. Comparison Analysis in IPA software was used to summarize multiple comparisons and visualized with pheatmap R package.

## Results

### Neonatal morphine exposure differentially altered the cellular composition in HIV, of the adolescent mouse midbrain

Single-cell RNA sequencing was performed on adolescent brains from wild-type (WT) and Tg26 mice exposed to morphine neonatally. Newborn WT and Tg26 mice were administered morphine at a concentration of 5 mg/kg/day for five days (Figure 1A). scRNA-seq libraries were generated from 8 mice midbrains. Midbrain tissue samples were sequenced using the 10X Chromium platform, which resulted in 77,073 high-quality cells retained after quality control. The retained cells were then computationally clustered in Seurat (11) and assigned to 13 specific cell types using the Sc-type pipelines (12) and its reference databases (Figure 1B). We then examined how NME alters the balance of cell types in the adolescent midbrain, specifically in the context of HIV. Across both groups, all neurons accounted for approximately 40% of the total cell counts. Oligodendrocytes and oligodendrocyte precursor cells (OPCs) constituted 25%, followed by astrocytes (15%), epithelial cells (6%), and microglia (5%), with the remaining cell types each representing less than 5% of the total cell count. A Wilcoxon rank-sum test was performed to determine if any cell type proportions differed significantly between the two conditions. This analysis revealed a significant decrease in cholinergic neurons (p= 0.036) from 18% to 12%, and a significant increase in GABAergic neurons (p= 0.036) from 9.5% to 17% following NME in the context HIV (Figure 1C). Among cell types comprising less than 5% population, tanycytic significantly decreased from 2.8% to 1.8% (p= 0.036) and dopaminergic neuron decreased from 0.7% to 0.3% (p= 0.036) (Figure 1C).

**Figure 1.**
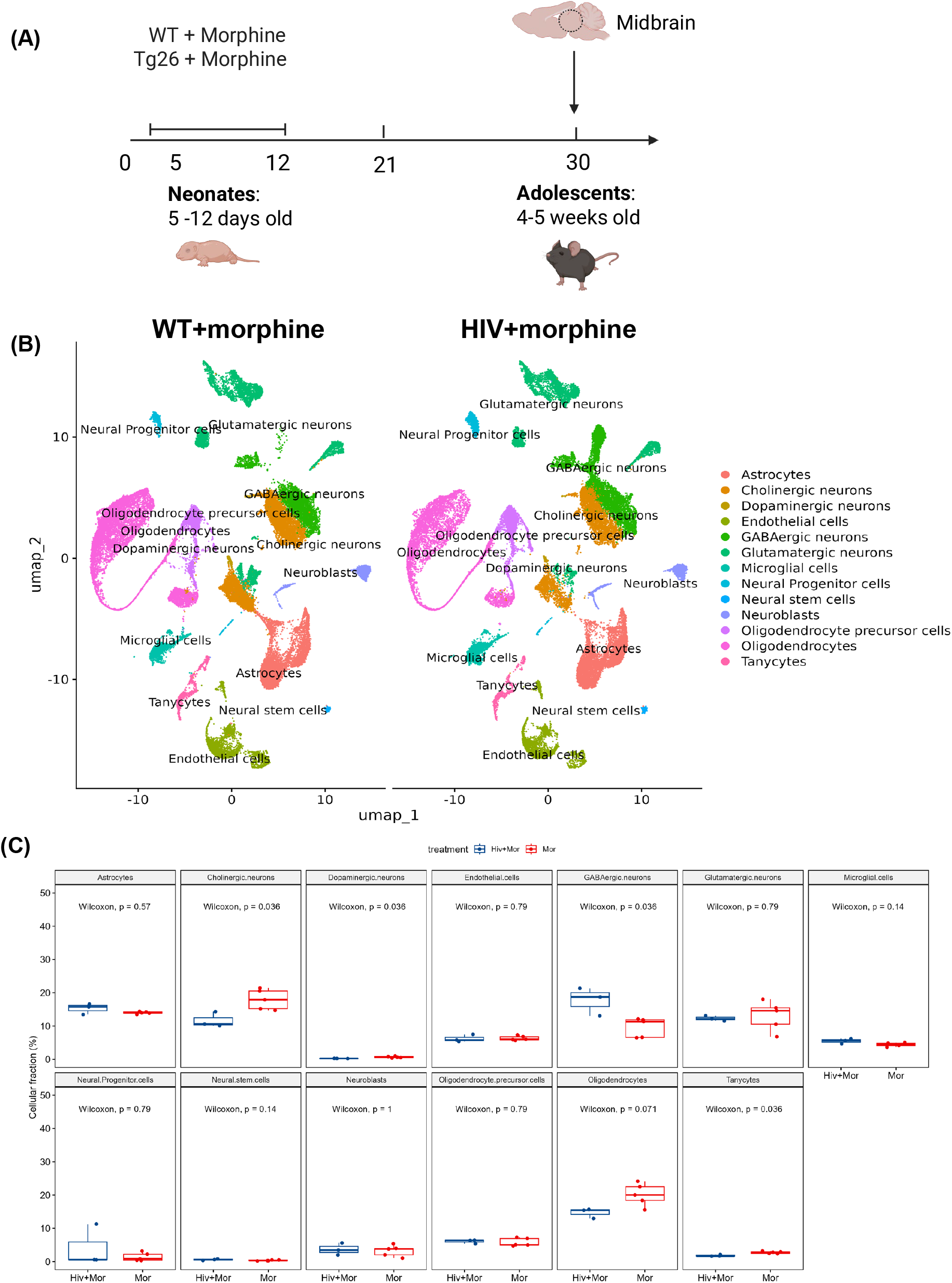
Morphine induced changes in cell type propotions in Tg 26 mice not observed in WT mice. A. Animal treatment design for 8 samples. WT+morphine (n=5), Tg26+morphine (n=3). B. Uniform Manifold Approximation and Projection (UMAP) plot showing the identified 13 major cell types, split by treatment groups for total of *n*□=□77,073 cells. C. Box plot of the 13 major cell types in each sample, split by treatment groups.

### Neonatal morphine exposure differentially altered the gene expression and pathways in HIV, in several cells of the adolescent midbrain

Next, we analyzed differentially expressed genes (DEGs) in microglia, glutamatergic neurons, GABAergic neurons, and cholinergic neurons to investigate transcriptional changes induced by neonatal morphine. Within the HIV genetic background, both the microglia and the neurons displayed significant transcriptional dysregulation following NME, as illustrated in the combined volcano plot (Figure 2). In microglia, the expression of genes for key neuromodulators (*Avp, Hcrt*, and *Pmch*) were upregulated, while genes related to immune defense (Map*3k6, Lgale3, Ccl3*, and *Il18rap*) were downregulated (Figure 2). Furthermore, QIAGEN IPA canonical pathway enrichment analysis showed that the opioid signaling pathway was upregulated in microglia, with expression of dynorphin (*Pdyn*) and κ-opioid receptor (*Oprk1*) significantly upregulated (Figure 3A). Pathogen-influenced signaling and immune response related pathway were concomitantly downregulated (Figure 3A). In cholinergic neurons, genes related to key neurohormone such as *Oxt* (oxytocin) and *Avp* (arginine vasopressin), as well as transcription factors such as *Fezf1* (FEZ Family Zinc Finger) and *Eomes* (Eomesodermin) were upregulated (Figure 2). Pathway analysis suggested that the function of mitochondrial is impaired in cholinergic neurons, as respiratory electron transport and oxidative phosphorylation were downregulated while mitochondrial dysfunction was increased (Figure 3B). In GABAergic neurons, genes with diverse functions were upregulated. These included transcription factors (*Lhx8*), genes encoding proteins involved in neuronal signaling (*Ppp1r1b*), ion transport (*Slc26a10*) and immune response (*Cd4*) (Figure 2). The pathway analysis in GABAergic neurons also demonstrated that several signaling pathways, including Camp mediated signaling, CREB signaling and G-protein signaling pathways, were upregulated (Figure 3C). In glutamatergic neurons, genes involved in homeostasis and behavior (*Avp, Pmch* and *Ot*), and homeobox transcription factors (*Dlx2* and *Dlx6*) were upregulated (Figure 2). Pathway analysis showed the opposite pattern in glutamatergic neurons compared to GABAergic neurons, with most neurotransmitter signaling being downregulated (Supplemental Figure 1).

**Figure 2.**
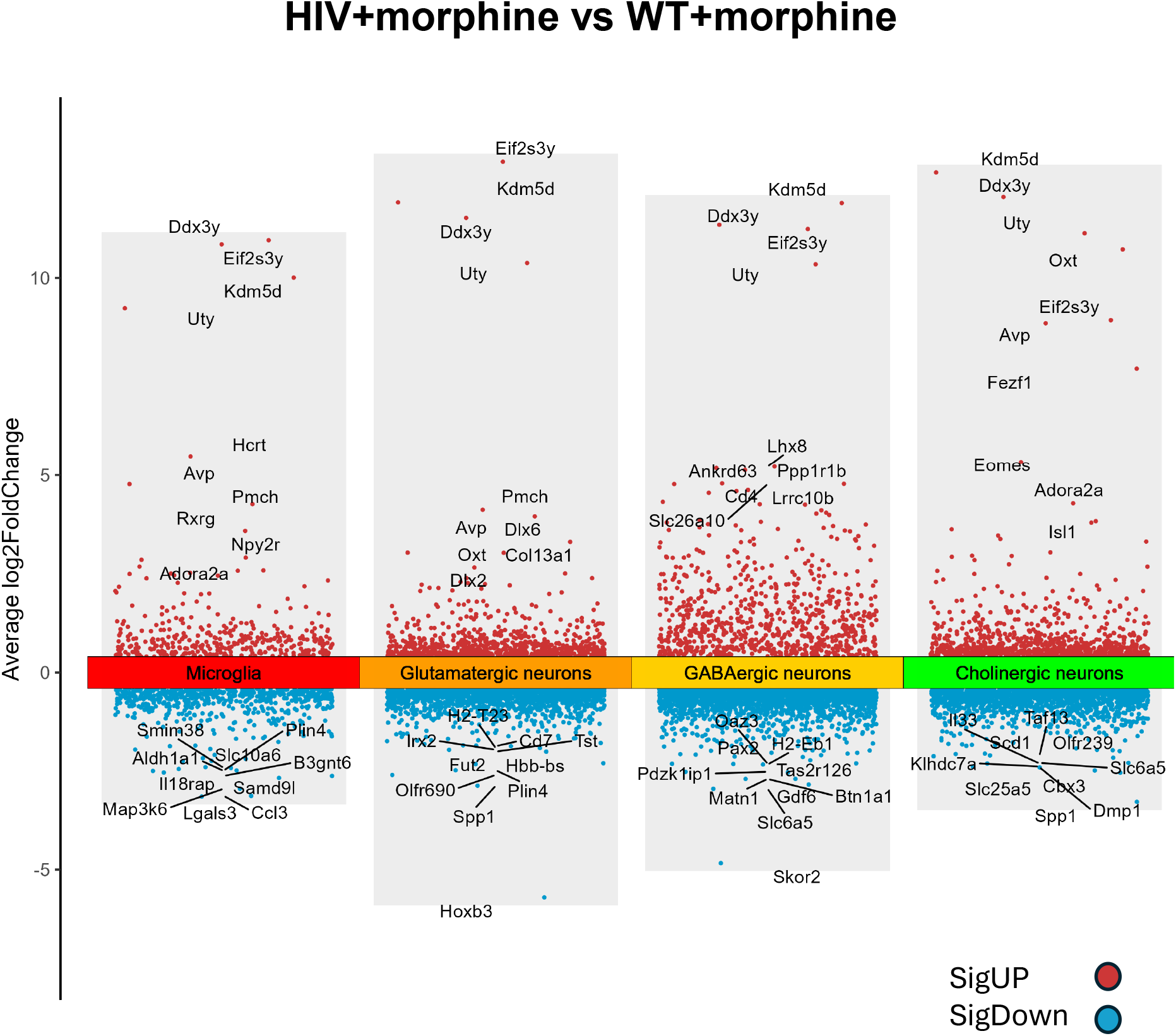
Differentially expressed genes (DEGs) under HIV+morphine vs WT+morphine. Combined volcano plots representation of differentially expressed genes (DEGs) across cell types. Genes with an absolute Log2 fold change (|Log2FC|) > 0.4 and an adjusted p-value < 0.01 were labelled statistically significant.

**Figure 3.**
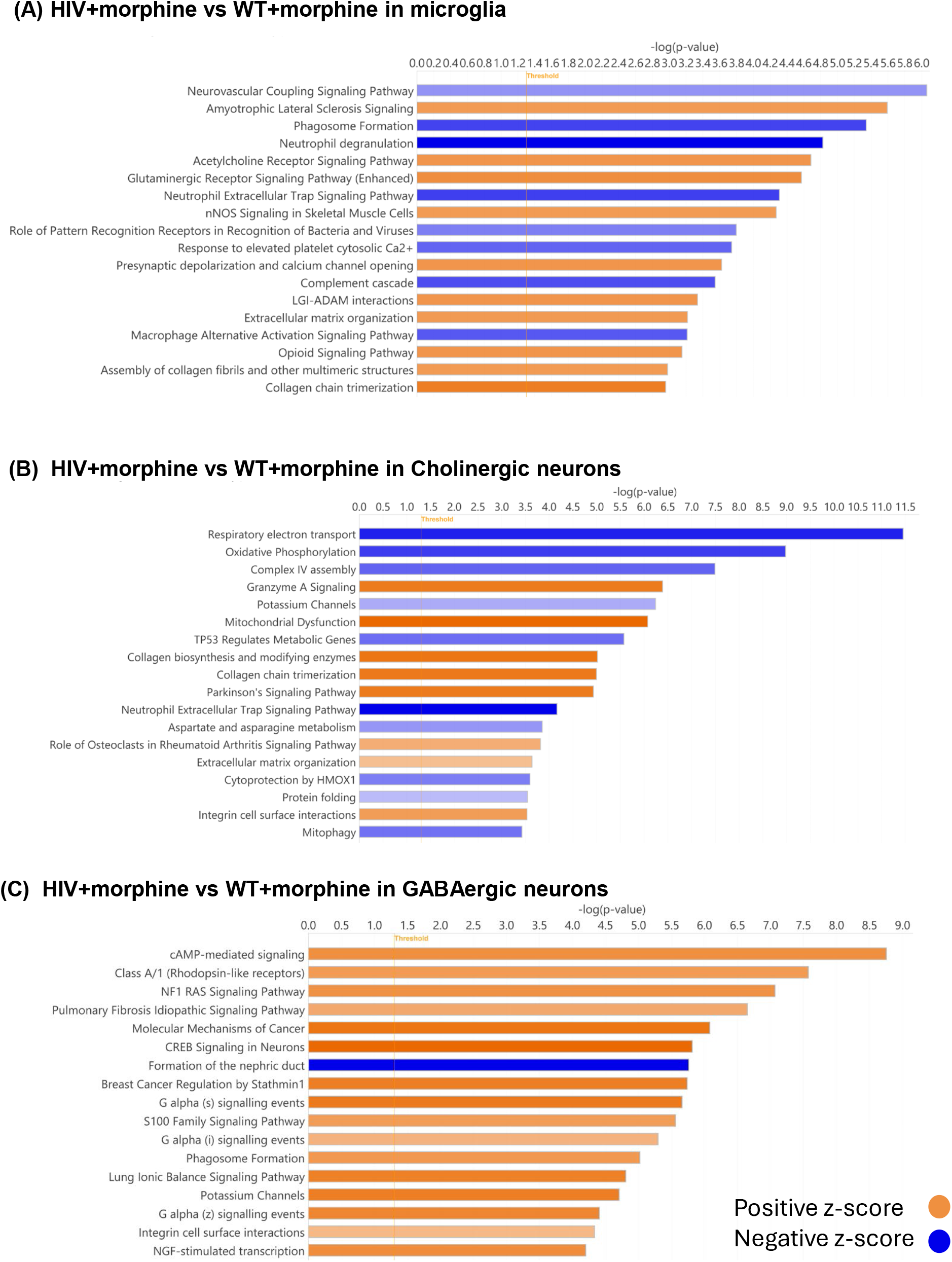
Canonical pathway changes induced by HIV background. Barplot of IPA canonical pathways in. (A) microglia under HIV+morphine vs WT+morphine, (B) cholinergic neurons under HIV+morphine vs WT+morphine, and (C) GABAergic neurons under HIV+morphine vs WT+morphine. Orange indicates positive z-scores, and blue indicates negative z-scores.

### Neonatal morphine exposure differentially induced behavior changes differently in HIV, including anxiety and fear in the adolescent midbrain

To further elucidate the downstream effect of the dysregulated gene expression, biological and disease functions were predicted using QIAGEN IPA with a specific focus on behavior changes. In cholinergic neurons, the observed gene expression profile was predicted to be associated with an induction of anxiety, emotional behavior, aversive behavior, and fear (Figure 4). Conversely, functions related to learning (including spatial learning and motor learning), cognition, and locomotion were predicted to be impaired (Figure 4). In GABAergic neurons, conditioning, learning and aversive behavior were predicted to be more likely to occur while aggressive behavior and thigmotaxis were less likely to occur (Figure 4). Conversely, in glutamatergic neurons, conditioning, and learning were less likely to occur, whereas thigmotaxis, emotional behavior and fear were predicted to be more likely to happen (Figure 4).

**Figure 4.**
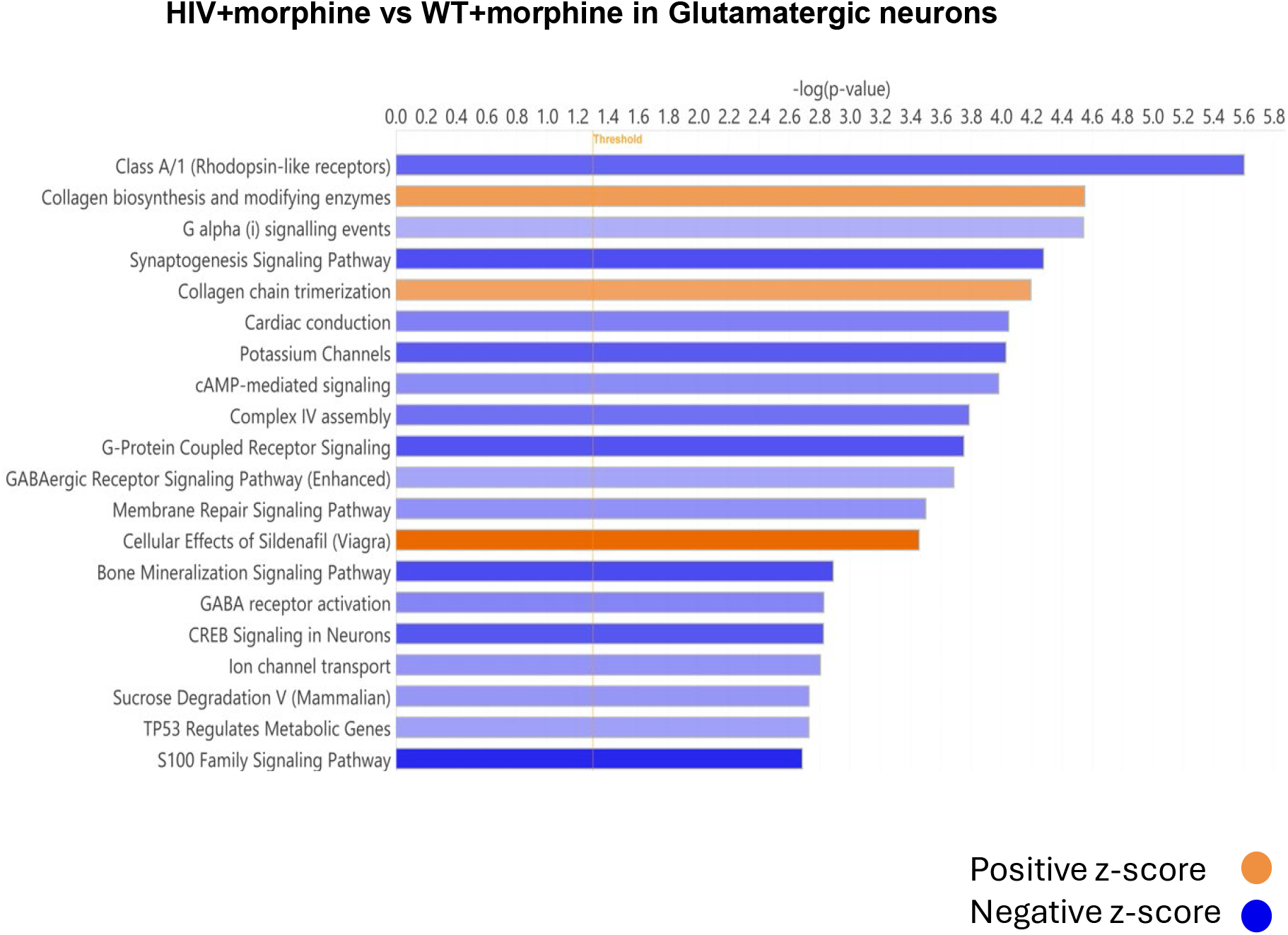
Behavior functions were altered by HIV background. Heatmap of top behavior functions for microglia, cholinergic neurons, and GABAergic neurons under HIV+morphine vs WT+morphine. Values are expressed in IPA z-scores.

## Discussion

This study demonstrates that brief NME, when superimposed on chronic HIV-related neuroinflammation, produces enduring and cell-type–specific transcriptional reprogramming in the adolescent midbrain. Using single-cell RNA sequencing, we show that HIV and morphine synergize to dysregulate microglial homeostasis and reshape neuronal signaling pathways governing anxiety, fear, and aversion. These findings provide mechanistic insight into why individuals with HIV who are exposed to opioids early in life may be at heightened risk for long-term affective vulnerabilities.

### HIV and morphine synergistically drive microglial reprogramming toward a maladaptive neuroimmune state

Microglia exhibited the most striking transcriptional perturbations. In Tg26 mice, NME induced an atypical neuropeptidergic microglial state characterized by ectopic upregulation of *Avp, Hcrt*, and *Pmch*, molecules typically restricted to hypothalamic and limbic neurons (13). Such ectopic neuromodulator expression suggests a shift toward a maladaptive, stress-responsive phenotype. Simultaneously, morphine-exposed Tg26 microglia displayed suppression of both inflammatory (*Map3k6, Lgals3, Ccl3*) and homeostatic (*P2ry12, Tmem119*) markers, indicating a loss of microglial identity consistent with chronic neuroimmune injury (14). The enrichment of dynorphin–κ-opioid receptor (*KOR*) signaling (*Pdyn, Oprk1*) within microglia is particularly notable. Dynorphin–KOR pathways are well-established mediators of dysphoria, negative affect, and stress-induced behavioral states (15). Their upregulation suggests that microglia, rather than neurons alone, may actively participate in shaping aversion-related circuits after early opioid exposure (16). Given that HIV infection independently promotes microglial activation and synaptic disruption (5–8), our findings indicate that NME imposes an additional and qualitatively distinct transcriptional burden, pushing microglia toward a hybrid neuroimmune–neuromodulatory phenotype poised to enhance anxiety susceptibility (17).

### Neuronal subtypes exhibit distinct, HIV-dependent alterations in anxiety- and aversion-related transcriptional programs

Across neuronal populations, we observed divergent adaptations that converge on circuits known to regulate emotional behavior. Cholinergic neurons showed robust upregulation of arousal-promoting neurohormones *Oxt* and *Avp*, as well as developmental transcription factors *Eomes* and *Fezf1*. IPA-based functional predictions linked these changes to increased anxiety, fear, and aversive behavioral outputs, with concomitant suppression of learning, locomotion, and spatial cognition. Such transcriptional patterns align with known cholinergic contributions to anxiety-like behavior and heightened sensory responsivity (18). GABAergic neurons displayed enhanced expression of aversion-associated transcripts, including *Lhx8, Ppp1r1b, Cd4*, and ion transporter *Slc26a10*. Upregulated cAMP, CREB, and G-protein coupled signaling pathways further point to heightened excitability or altered inhibitory tone. CREB-mediated pathways in GABAergic circuits are strongly implicated in negative effects, stress conditioning, and opioid-induced aversion. Glutamatergic neurons shifted in the opposite direction, with downregulation of multiple neurotransmitters signaling pathways and upregulation of behavioral genes associated with thigmotaxis and fear (e.g., *Avp, Pmch, Dlx* family transcription factors). Collectively, these adaptations imply a dampening of excitatory homeostasis and a parallel enhancement of environmental avoidance behaviors. The net effect across neuronal subtypes is consistent with the well-described imbalance between excitatory and inhibitory signaling observed in models of anxiety and HIV-associated neurocognitive disorders. Importantly, these transcriptional signatures emerge weeks after cessation of morphine exposure, demonstrating that early-life opioid exposure produces durable programming of emotional circuits, especially in an HIV-inflamed environment.

### Integration with prior literature and conceptual model

Previous work has shown that chronic or escalating opioid exposure interacts with HIV proteins such as Tat or gp120 to amplify neuroinflammation, synaptic injury, and behavioral impairments (5–8). However, the effects of brief neonatal exposure, a clinically relevant scenario for infants treated with opioids for neonatal abstinence syndrome or those exposed perinatally have remained largely unexplored. Our data suggest that even short-term morphine exposure during a critical neurodevelopmental window may alter microglial maturation trajectories and disrupt the assembly of cholinergic, GABAergic, and glutamatergic circuits. The Tg26 model, which recapitulates chronic viral protein–mediated immune activation, appears to sensitize the developing brain to opioid-induced transcriptional injury. We propose a two-hit mechanism in which: HIV-driven neuroinflammation primes microglia and neurons, reducing their resilience and altering developmental trajectories, and Neonatal morphine imposes an additional neuromodulatory stressor, triggering maladaptive dynorphin– KOR signaling, suppression of metabolic pathways, and long-term reprogramming of affective circuitry. The convergence of these processes on pathways governing fear, aversion, and anxiety is consistent with clinical observations that opioid-exposed and HIV-exposed youth face heightened risk for emotional dysregulation, stress hypersensitivity, and substance-use vulnerability later in life (1–4).

### Limitations and future directions

Although the single-cell approach provides high-resolution insight into cell states, several limitations should be considered. First, we focused on the midbrain; additional regions such as the amygdala, hippocampus, or prefrontal cortex may reveal complementary or distinct adaptations. Second, transcriptional states do not fully capture functional consequences; future studies integrating electrophysiology, in vivo imaging, and behavioral assays will be essential. Third, although the Tg26 model recapitulates chronic HIV protein exposure, it does not fully model viral replication or ART effects. Studies using EcoHIV or humanized mouse models would provide deeper translational insight. Nevertheless, our identification of dynorphin–KOR signaling and microglial neuromodulator dysregulation as core nodes provides compelling targets for mechanistic validation and potential therapeutic intervention.

## Conclusions

In summary, we show that neonatal morphine exposure in an HIV-inflamed brain produces persistent and cell-type–specific transcriptional reprogramming across microglia and key neuronal subtypes. These changes converge on pathways promoting anxiety, fear, aversion, and impaired learning, highlighting a microglia–neuron interaction network that may underlie long-term neuropsychiatric vulnerability in opioid- and HIV-exposed youth. By resolving these early transcriptional alterations at single-cell resolution, our study opens new avenues for identifying developmental windows and molecular pathways that could be targeted to mitigate neurobehavioral risk in this growing and vulnerable population.

## Figure Legends

**Supplemental Figure 1.**
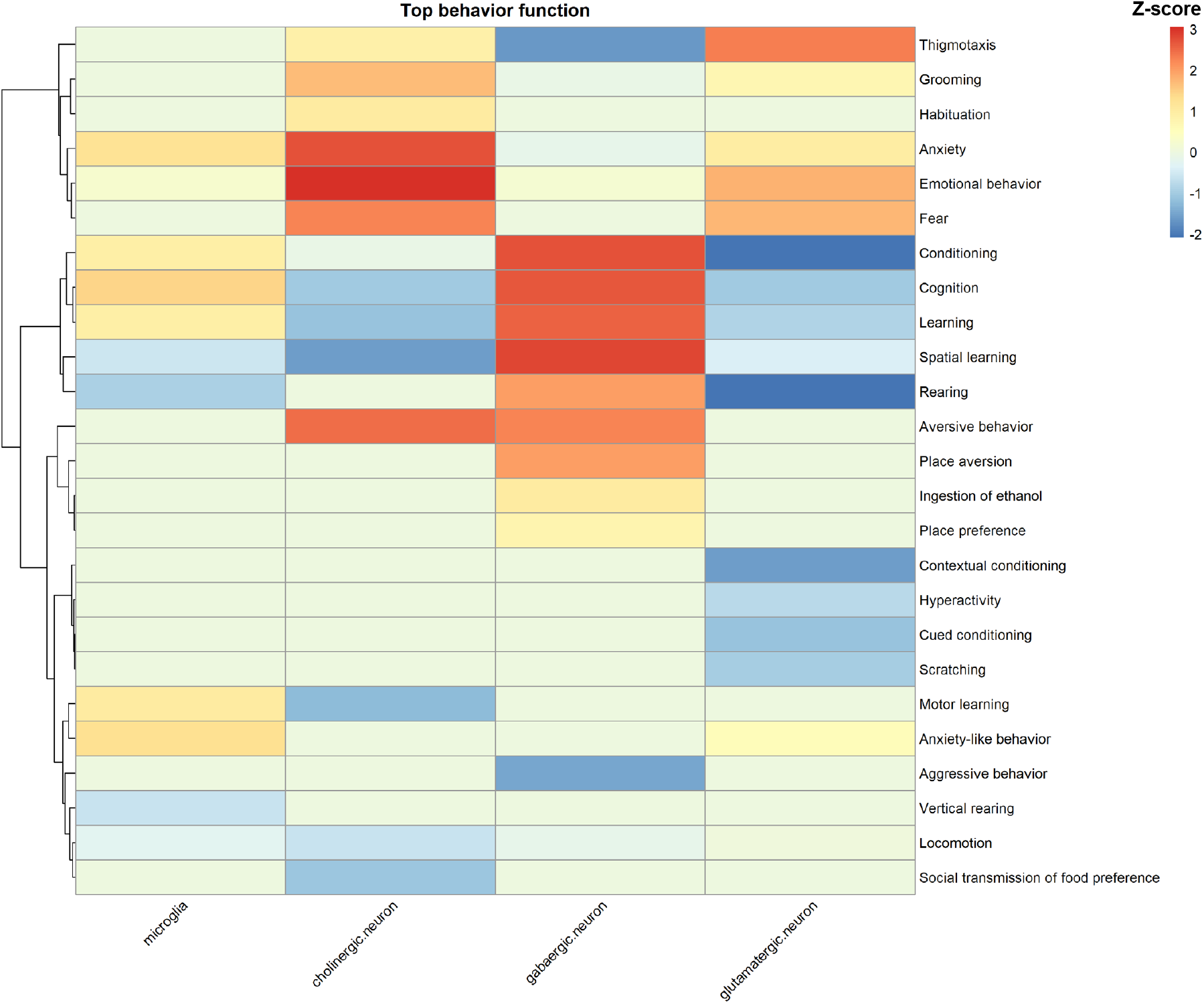
Canonical pathway changes induced by HIV background. Barplot of IPA canonical pathways in glutamatergic nueonrs under HIV+morphine vs WT+morphine,

